# Hidden Bias: GPCR Polymorphisms and the Genomic Landscape of β-Arrestin Signaling

**DOI:** 10.1101/2025.06.03.657708

**Authors:** Yang Zhou, Alp Kuzey Sozat, William C. Wetsel, Lauren Slosky, Joshua Gross, Lawrence S. Barak

## Abstract

Clinically, no human diseases are currently diagnosed as being directly driven by β- arrestins. Nonetheless, there is growing interest in developing β-arrestin–biased drugs, due to the key regulatory role of β-arrestins in modulating the cellular signaling of most G protein–coupled receptors (GPCRs). In cell-based studies of rhodopsin-family GPCRs, mutations in a conserved proline–hydrophobic (ProH) motif within the second intracellular loop (ICL2) have been shown to alter β-arrestin signaling bias. The clinical relevance of such mutations in humans is unknown. However, if naturally occurring single nucleotide polymorphisms (SNPs) affecting this ProH motif exist across the GPCR family, then cross-referencing these genetic variants with large-scale population sequencing and epidemiologic data could reveal potential roles for β-arrestin signaling in human health. In this report, we identify SNPs in human GPCRs that correspond to ProH substitutions, and we show that, in the neurotensin receptor NTSR1, these variants can indeed shift signaling bias. We also estimate a lower-bound population frequency for such SNPs, suggesting that although rare for any given receptor, their cumulative prevalence across the GPCR superfamily may be large enough to impact phenotypic variation. Together with emerging data from biased ligands, our findings support the idea that genetic variations in β-arrestin signaling could represent a meaningful source of therapeutic relevance.

**Significance statement:** Mutations altering G protein-coupled receptor (GPCR) signaling can affect health. We have identified a specific amino acid determinant in intracellular loop (ICL2) of human rhodopsin family GPCRs that influences β-arrestin signaling. Although the medical consequences of this determinant remain unclear, we show that single nucleotide polymorphisms (SNPs) affecting the determinant routinely occur. By studying neurotensin receptor NTSR1, we confirm that the SNP alters β-arrestin signaling bias. While individually rare per receptor, these SNPs may collectively contribute to β-arrestin-related phenotypic changes in the human population. Our findings, combined with research on biased drugs, support the idea that β-arrestin signaling might serve as a useful therapeutic target, opening new possibilities for precision medicine.

## Introduction

G protein coupled receptors (GPCRs) are exceptionally good drug targets for treating a wide range of diseases. Many drugs such as adrenergic receptor antagonists typically affect GPCR regulation in an unbiased manner by similarly modifying activity of G protein and β-arrestin (β-arrestin) signaling pathways (1, 2). GPCRs control many areas of human physiology, signaling through four families of heterotrimeric G proteins and two ubiquitously expressed non-visual beta-arrestins β-arrestins(3). Together the G proteins and β-arrestins form the principal intracellular components for GPCR signaling scaffolds (4). It is well established that receptor/G protein complexes play an important role in the translation of chemical signals to physiological function, but only relatively recently has a non-desensitization, direct, independent role for β-arrestin scaffolds been proposed in GPCR signaling biochemistry (5). We have previously shown for representative rhodopsin family GPCRs a proline-hydrophobic (ProH) motif in receptor intracellular loop 2 (ICL2) affects β-arrestin interaction with the receptor core and can change the relative extent of G protein/β-arrestin activation(6). The (ProH) motif is +6 downstream of the highly conserved third transmembrane (D/E)RY motif, itself an important determinant for ligand-mediated receptor activation(7). We also know that the (ProH) motif signaling changes vary with the type of amino acid mutation(6), but we do not know if such effects on biochemistry translate into variations in physiology. Our initial goal is two-fold: first to determine that the existence of (ProH) motif SNPs is generalizable, and second to assess whether cellular changes in β-arrestin signaling occur as a result. It is well established that GPCR variants result in human disease(8). We believe the ProH motif will be important clinically because of its regulatory position and conservation in rhodopsin family GPCRs. Understanding how variations in the ProH motif modify function addresses fundamental questions in β-arrestin signaling biology and will improve strategies for developing biased drugs(1).

## Results

We previously determined from the NCBI (National Center for Biotechnology Information) protein database that among the rhodopsin family receptors amino acids P and A occupy the first ProH site with 96% frequency(6) (**Figure 1A**), and that the chemokine receptor subfamily is where substitution of proline by alanine is most commonly observed(6). Notably, AVPR2, the vasopressin receptor gene that regulates renal water balance, has such a missense mutation, rs782571215 (C>G). The substitution occurred in 2 individuals homozygous for the G allele at variant frequencies 1.1 × 10^−5^ and 2.3 × 10^−5^ per respectively the GnomAD exome and Exome Aggregation Consortium databases. Clinical phenotypes were not listed. A proximate -7 SNP, rs104894756 for AVPR2, chrX:153905916, frequency:1/176496, bordering ICL2 at the AVPR2 DR(H/Y) motif arginine, however, causes β-arrestin-dependent diabetes insipidus(9). (Unless otherwise noted missense SNP data are referenced from NCBI dbSNP, missense variant, September 21, 2022, build 156 and study: GnomAD exome.) This suggests ICL2, particularly at the ProH may be a hot spot for SNPs affecting β-arrestin signaling bias. We, therefore, wondered how easily other receptors with ProH motif missense could be found, and, if found, what if any effect the mutation would have on intracellular signaling.

**Figure 1.**
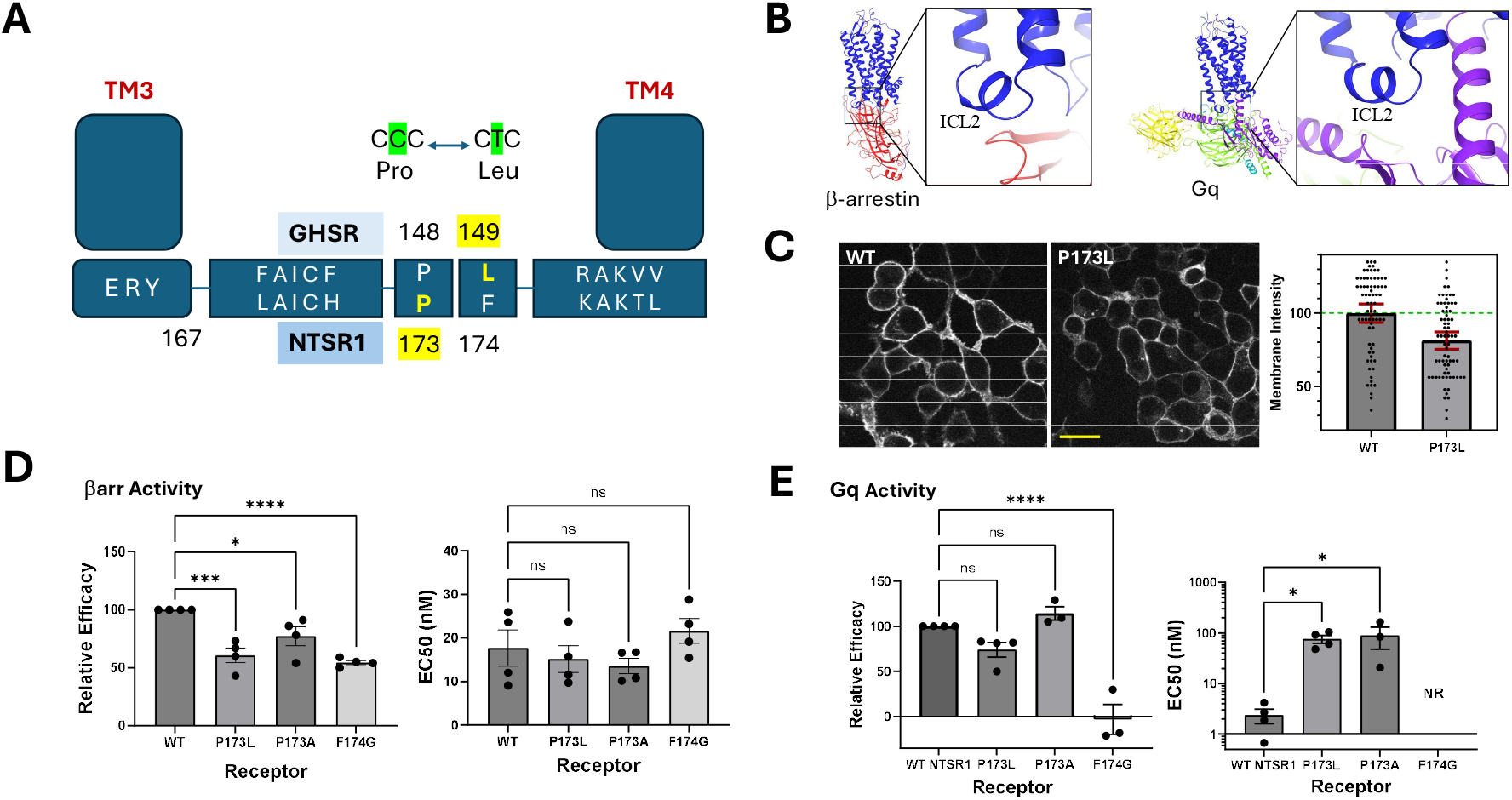
NTSR1 Mutation and Signaling. (**A**) Cartoon depiction of the linear structure following the ERY motif of the NTSR1 at the C>T substitution P173 and the GHSR at the T>C substitution in L149P. (**B**) Cryo-EM structures of NTSR1 (blue) bound to β-arrestin2 (left, red, PDB ID 6PWC) or Gq (right, purple, PDB ID 8FMZ) in the location of the receptor ICL2. (**C**) Immunofluorescence micrographs (images on left) of NTSR WT and P173L mCherry-GFP fusion protein variants, relative intensities with the local background subtracted were determined using ImageJ and were expressed (bar graph on right) as mean ± 95% confidence interval, WT (1.0 ± 0.06) and P173L (0.81 ± 0.06, p<0.0001) (n = 72 sampled sites each from 6 image fields obtained from two independent transfections, yellow size bar = 20 microns). (**D**) For β-arrestin activity (Emax, EC50) of the receptors relative to WT NTSR1 (100%, 18 ± 5 nM) are: P173L (61± 7%, *** p = 0.0002), (15 ± 4 nM, p = 0.58), P173A (77 ± 9%, * p = 0.011), (14 ± 2 nM, p =0.37) and F174G (55 ± 2%, p < 0.0001), (22 ± 3 nM, p = 0.38), (N=4). (**E**) For Gq activity (Emax, EC50) of the receptors relative to WT NTSR1(100%, (2.4 ± 1) nM) are: P173L (74 ± 8%, p=0.051), (76 ±13 nM, * p = 0.03), P173A (114 ± 7%, p = 0.28), (89 ± 41 nM, * p = 0.02)) and F174G ((−3 ± 17, **** p < 0.0001) %, NR = not reported), (N=4 for WT and P173L and 3 otherwise). For (D-E) data are presented as mean ± sem and were analyzed in GraphPad Prism version 10 (Dotmatics, Boston MA) using one way ANOVA with Fishers Uncorrected Least Significant Difference (LSD) post-hoc test.

### Search for ICL2 ProH SNPs in representative GPCRs

We began our search with examination of the NCBI SNP database for the human growth hormone secretagogue receptor (GHSR) and the human neurotensin 1 receptor (NTSR1) because we already had developed signaling biased drugs for both (10-12). We did not have to look further. We found a naturally occurring ICL2 ProH variant for GHSR, a hydrophobic substitution GHSR^L149P^ (SNP rs1390872592, chr3:172447968, frequency: 1/250994), and subsequently one for NTSR1, a proline residue substitution NTSR1^P173L^ (SNP rs1383266510, NTSR1, chr20:62709725, frequency: 3/264690 and study: TOPMED, frequency: 1/140272) (**Figure 1A**). Herein we briefly characterize signaling for the rs1383266510 variant NTSR1^P173L^ and compare it to the wild-type NTSR1 behavior. Elsewhere in a companion manuscript (Balfe et al., 2025, *BioRxiv*), we provide a detailed explanation of the molecular mechanisms underlying the signaling bias of the natural variant GHSR^L149P^.

### Prevalence of β-arrestin regulatory mutations in the ProH motif

We developed a general formula to estimate the minimum number of expressed variant carriers N_E_ for a population N_p_. N_E_ should be proportional to the number of independent receptor sites (N_s_) with SNPs, the number of receptors (N_G_), and the mutation frequencies M_F_ because each SNP is relatively rare. Thus, N_E_ = N_s_ x N_G_ x M_F_ x N_p_. With SNP frequencies for GHSR and NTSR1 of order N_M_ = 0.5-1 x10^−5^ per person, the number of non-odorant rhodopsin family GPCRs of order N_G_ = 2.5 x10^2^, and the number of potential sites for mutagenesis affecting the ProH determinant minimally N_s_ = 2, then N_E_ = (1/200 – 1/400) N_p_.

### The NTSR1 P173L mutant has reduced β-arrestin2 recruitment

NTSR1 signals predominantly through receptor/G_q_ complexes and strongly recruits β-arrestins following stimulation with neurotensin (NT) (**Figure 1B**) (12). NTSR1 signaling produces neuroleptic effects, regulates energy utilization and fat metabolism, and is growth promoting (12). β-arrestins bind, desensitize, and traffic NTSR1s by scaffolding clathrin adapters and numerous other downstream signaling proteins (4). Both the NTSR1 and NTSR1^P143L^ express at the plasma membrane when transfected into HEK293T cells (**Figure 1C**). We used a BRET proximity assay for NTSR1/ β-arrestin2 over a NT range of 5 ½ decades to determine their response maxima as a measure of relative β-arrestin activity (**Figure 1D**). For added reference, we evaluated NTSR1^P173A^, and NTSR1^F174G^ variants, artificial mutations that have been shown previously for the GHSR1a to alter β-arrestin or Gq activity (**Figure 1D**) (13). A significant difference for interaction with β-arrestin2 was observed between the wild-type E_max_ (100% ± 0%) and P173L (62% ± 12 %) variant. In this assay, a reduction in Emax may reflect a reduction in the number of NTSR1-β-arrestin2 complexes or a change in complex conformation that increases the distance between BRET donor and acceptor. No significant difference was observed between the wild-type and variant potency (EC_50_).

### P173L mutant has reduced Gα_q_ activation

Gα_q_ activation was measured over a similar NT range using a BRET2 proximity assay TRUPATH (14) (**Figure 1E**). We observed a significant 25% reduction in Gα_q_ activation efficacy between WT and P173L (100% versus 74 % ± 8% respectively). Most importantly, compared to WT receptor, the NTSR1^P173L^ as well as the other mutants had more than 25-fold reduced potency. This large reduction in Gα_q_ signaling potency indicates that the mutation has the potential to physiologically bias NTSR1^P173L^ signaling relative to the WT receptor, reflected by the corresponding bias factor of β=1.3 (15), a β= 0 indicating no bias. Surprisingly, the NTSR1^F174G^ variant was completely unable to activate Gα_q_ (−3% ± 17%) over the range of NT tested.

## Discussion and Summary

Our SNP variant data is limited to a restricted population that has undergone genomic sequencing. Nonetheless, two arbitrarily selected GPCRs with ProH variants that, taken together with the natural variation at the proline in chemokine receptors and the AVPR2 (rs782571215, (P144 >A), chrX:153905936, frequency: 2/176730, and frequency: 2/86820, study: ExAC) suggest that mutations in the ProH motif could be common throughout the rhodopsin family of GPCRs. Thus, as population genotyping becomes more widespread, the chances of establishing clinical correlations in areas like diabetes insipidus (AVPR2), heart disease, and obesity, for example with ProH motif-containing receptors like the β1-AR rs1847533244, (P163>L), chr10:114044620, frequency: 0/10680, study: ALFA) or orexin receptor HCRTR1 rs1004014305 (P151>T, HCRTR1, chr1:31620915); rs200634924 (P151>L, chr1:31620916) should increase by exploiting mutations that change β-arrestin/G protein signaling bias. Our lower bound prevalence estimate of 2500-5000 per million individuals with these SNPs together with the further development of biased drugs should eventually enable defining the degree to which β-arrestin can affect pathophysiology, general health, and response to disease treatment in human populations.

## Materials and Methods

### Plasmids

Receptor variants fused to the N-terminus of mCherry fluorescent protein using a 14 amino acid polylinker were synthesized by Genscript (Piscataway NJ). The TRUPATH assay was a gift from Bryan Roth (Addgene kit #1000000163).

### Chemicals

All chemicals were obtained from MilliporeSigma (St. Louis, MO, USA) unless otherwise noted. Coelenterazine h and 400a were obtained from NanoLight (Phoenix, AZ, USA, Cat#301 and Cat#340). For receptor signaling assessments, NT (Sigma, product #N6383) was maintained as a glycerol stock.

### Cell lines

HEK293T/17 (Cat# CRL-11268, RRID:CVCL_1926) cells were obtained from the American Type Culture Collection (ATCC, Manassas, VA.

### Cell Assays

**β-arrestin** (β-arrestin BRET) – Cells (HEK293T) were calcium phosphate transfected using 1.5 μg of an mVenus β-arrestin2 plasmid and 100 ng of 3xHA-hNTSR1-Rluc8 plasmid from WT, P173L, P173A, or F174G receptor. **G Protein** (TRUPATH assay) **-** Cells were transfected using 1.6 μg pcDNA3.1, 100 ng of each of three G protein subunit plasmids (Gq alpha-RLuc8, Gβ3, and Gγ9-GFP2), and 100 ng 3xHA-hNTSR1 plasmid cDNA from WT, P173L, P173A, or F174G receptor. Transfected cells (60,000) were split into 96 well poly-D-lysine treated plates (Corning Life Sciences, Kennebunk, ME, #3610) in opti-MEM with 2% FBS and 1x antibiotic antimycotic solution for incubation overnight at 37°C in 5% CO2. BRET acceptor/donor values of (535-40 nm)/(475-30 nm) or (515-30 nm)/(410-80 nm), for β-arrestin BRET or TRUPATH assay respectively, were measured using a ClarioStar microplate reader. BRET ratios were calculated by dividing the raw acceptor values by the raw donor values and net BRET ratios by subtracting the control BRET ratio (no NT) averaged over 3 replicates from the NT-treated well BRET ratio

### Immunofluorescence

*HEK293*T cells in a 35 mm MatTek (Ashland, MA, CAT# P35GC-1.5-14-C) optical dish were with transfected with 1.5 µg receptor and 1.0 µg pcDNA3.1 using lipofectamine 3000 (Thermo Scientific, Waltham, MA, CAT# L3000015) per the manufacturer’s recommendations and imaged live the next day in optimum buffered with 10 mM Hepes at 568 nm excitation with a rhodamine bandpass filter using a 40X 1.3 n.a. oil objective.

## Acknowledgements

We would like to thank Mohanraj Krishnan, Assistant Professor of Biobehavioral Health, Department Pennsylvania State University State College, Pennsylvania for reviewing the manuscript. The authors have no conflicts of interest This grant was partially funded by a subaward funded by the National Institute on Drug Abuse (NIDA) under grant number #61174-13926, administered through Duke University.

